# Impact of Unc-51 Like Kinase 4 (*ULK4*) on the Reactivity of the Extended Reward System in Response to Conditioned Stimuli

**DOI:** 10.1101/2024.01.15.575326

**Authors:** Jens Treutlein, Simone Löhlein, Karolin E. Einenkel, Esther K. Diekhof, Oliver Gruber

## Abstract

**Objectives:** *ULK4* is an established candidate gene for mental disorders and antipsychotic treatment response and codes for a serine/threonine kinase that regulates the neural stem cell pool and controls cortex development. We investigated the effects of functional genetic variation at the *ULK4* locus on the human extended dopaminergic reward system using functional magnetic resonance imaging (fMRI) during performance of a well-established reward paradigm.

**Methods:** 234 study participants with functional neuroimaging (fMRI) data of the extended reward system and with *ULK4* genotype data were included in this study. Effects of genetic variation in the *ULK4* gene on reward system functioning were determined using the Desire-Reason-Dilemma (DRD) paradigm which allows to assess brain activation in response to conditioned reward stimuli (Diekhof et al. 2010).

**Results:** Among common missense variants of the *ULK4* gene, variant prioritization revealed strongest functional signatures for variant rs17215589, coding for amino acid exchange Ala715Thr. For rs17215589 minor allele carriers, we detected increased activation responses to conditioned reward stimuli in the ventral tegmental area, the nucleus accumbens and several cortical brain regions of the extended reward system.

**Conclusions:** Our findings provide further evidence in humans that genetic variation in *ULK4* may increase the vulnerability to mental disorders by modulating the function of the extended reward system. Future studies are needed to confirm the functional modulation of the extended reward system by ULK4 and to specify the role of this mechanism in the pathogenesis of psychiatric disorders.

## Introduction

The reward system in the brain is based on a neural circuitry including regions of dopaminergic neurotransmission, in particular the ventral tegmental area (VTA) and the ventral striatum (VS), which act in concert with more distant brain regions^1, 2^, forming the extended reward system. Evidence for disturbances in these regions in psychiatric disorders has been provided by functional magnetic resonance imaging (fMRI) studies using reward-specific paradigms^3-7^.

With respect to studies in imaging genetics, to date most reported genetic associations with reward system functioning in humans derive from the classical dopaminergic candidate genes from early studies in model organisms^8-10^. Firing of dopaminergic VTA neurons, which form synapses with neurons in the nucleus accumbens, was found to be crucial for the encoding of reward information^1^. Therefore, it is not surprising that most reported genetic associations in humans refer to the dopamine system. After release, dopamine is taken up into the presynapse of the VTA neuron by the dopamine transporter (DAT1). In line with this function, genetic variation of DAT1 has also been identified to modulate the reward system^11, 12^. After diffusion across the synaptic cleft, dopamine binds to postsynaptic receptors in the nucleus accumbens neuron, and to presynaptic autoreceptors in the VTA neuron. Consistent with this knowledge it was shown that genetic variation at the dopamine D2 receptor modulates striatal fMRI responses during a monetary reward paradigm^13^. Activated dopamine receptors in turn influence intracellular signalling cascades to cause gene expression differences via transcription factors, e.g. CREB1. As can be expected, genetic variation of *CREB1* was shown to influence reward system activity as well^14^.

Until now, only few other candidates whose relationship to dopaminergic neurotransmission is less obvious have been reported to influence the reward system as well. First, *MAD1L1*, whose primary role is cell cycle control, has initially been identified in a genome-wide association study (GWAS) for bipolar disorder, and subsequently was shown to influence reward system reactivity by imaging genetics analyses^15^. Another gene that is only indirectly linked to dopaminergic neurotransmission is vacuolar protein sorting-associated protein 4a (*VPS4A*), which was implicated in a GWAS for reward anticipation and functions in intracellular protein transport^16^.

A further possibly indirect dopaminergic candidate gene is Unc-51 Like Kinase 4 (*ULK4*), which was found to be required for axonal elongation in the model organism *Caenorhabditis elegans*^17^. It was observed that *ULK4* is substantially upregulated by treatment with retinoic acid, which leads to a dopaminergic-like phenotype^18^, in a human neuroblastoma cell line^19^.

For schizophrenia and affective disorders, which are both related to dopaminergic dysfunction^20^, *ULK4* constitutes an established susceptibility gene^19, 21^. Copy number variation data from the International Schizophrenia Consortium showed that deletions of *ULK4* were present in schizophrenia patients, but not in controls. Similar enrichment of *ULK4* in schizophrenia and bipolar disorder was detected in Icelandic cases by deCODE^19^. Furthermore, SNPs in *ULK4* were reported to be associated with antipsychotic treatment response^22^.

The triangulation between *ULK4*, psychiatric disorders, and dopaminergic neurotransmission motivated us to investigate the effects of functional genetic variation of *ULK4* on the neurofunctional level. To this end, a homogenous group of healthy adults underwent fMRI. All participants performed a specific reward paradigm, the ‘Desire-Reason Dilemma’ (DRD) paradigm^23, 24^, to investigate possible gene effects on the mesolimbic reward system and other reward-related brain regions. We hypothesized that carriers of functional allelic variants of this gene would show differences in the reactivity of the human extended reward system in response to conditioned reward stimuli.

## Methods

### Subjects

Participants of the Genomic Imaging Goettingen (GIG) study (N=299) were recruited by advertisements in intern online student networks and local newsletters in the Georg-August-University Goettingen and the University Medical Center Goettingen. Healthy young adults aged 18–31 years were included.

Exclusion criteria were past or present psychiatric disorders according to ICD-10, a positive family history of psychiatric disorders, substance abuse during the last month, cannabis abuse during the last 2 weeks, mental retardation, dementia, neurological or metabolic diseases, and pregnancy in women. All participants were of European ancestry. After exclusion of 64 individuals due to imaging data quality control, genotyping data were available for N=235 participants. For one subject the genotype was missing for *ULK4* variant rs17215589, leaving N=234 individuals for imaging genetic analyses.

The study was carried out in accordance with the Declaration of Helsinki and was approved by the local ethics committees, of the Medical Faculty of Göttingen University (number 14/3/09, date 02.07.2009) and of the Medical Faculty of Heidelberg University (number S-123/2016, date 09.03.2016). All participants provided written informed consent.

### Experimental Procedure / Desire-Reason-Dilemma Paradigm

Initially, participants underwent an operant conditioning task. Eight differently colored squares were presented as stimuli on a monitor in a shuffled mode. Subjects were instructed to respond to each of the stimuli by button press with their right hand. Button choice was free and subjects were encouraged to explore the stimulus-response-reward contingencies. By doing so, subjects were conditioned to associate two colors (red and green) with an immediate reward (bonus of +10 points), while the other six colors were associated with a neutral outcome. The goal of this operant conditioning task was to establish stimulus-response-reward contingencies for the next phase of the experiment.

Subsequently, subjects were familiarized with the actual experimental task, the DRD paradigm, a delayed matching to sample task. Subjects had to perform blocks of four or eight trials. At the beginning of every block, subjects were shown two targets (two different neutral colors, not the previously conditioned colors red and green). In the following, four or eight colored squares were presented one after another. To achieve the superordinate goal (50 points at the end of each block), subjects had to accept the two target colors shown at the beginning and to reject non-target colors by button press. Two different types of blocks had to be performed. For the present project, only the first type of blocks, the ‘Desire Context’ (DC) was relevant. In the DC, subjects are allowed to accept the previously conditioned reward stimuli in addition to the two target colors in order to win bonus points^23, 24^.

### fMRI Data Acquisition, Preprocessing and Analysis

fMRI was performed on a 3-Tesla Magnetom TIM Trio Siemens scanner (Siemens Healthcare, Erlangen, Germany) equipped with a standard eight-channel phased-array head coil. First, a T1-weighted anatomical data set with 1 mm isotropic resolution was acquired. Parallel to the anterior commissure– posterior commissure line, thirty-one axial slices were acquired in ascending direction for fMRI (slice thickness = 3 mm; interslice gap = 0.6 mm) using a gradient-echo echo-planar imaging sequence (echo time 33 ms, flip angle 70°; field-of-view 192 mm, interscan repetition time 1900 ms).

In two functional runs, 185 volumes each were acquired. Subjects responded via button presses on a fiber optic computer response device (Current Designs, Philadelphia, Pennsylvania, USA), and stimuli were viewed through goggles (Resonance Technology, Northridge, California, USA). Presentation Software (Neurobehavioral Systems, Albany, California, USA) was used to present the stimuli in the scanner.

Functional images were preprocessed and analyzed with SPM12 (Statistical Parametric Mapping; www.fil.ion.ucl.ac.uk/spm/software/spm12/) using a general linear model. The study design was event-related and only correctly answered trials were included in the analysis.

Linear t-contrasts were defined to assess brain activation effects elicited specifically by the conditioned reward stimuli as compared to non-rewarded stimuli. These single-subject contrast images were taken to the second level to assess genotype effects using two sample t-tests contrasting minor allele carriers with major allele homozygotes. Whole-brain genotype group effects were searched for using P<0.005, uncorrected, as a search criterion for further statistical evaluation.

### Genotyping, Variant Selection and Effect Prediction

Saliva was collected using Oragene DNA devices (DNA Genotek, Ottawa, Ontario, Canada), and DNA was isolated with standardized protocols. Genotyping was performed using Illumina OmniExpress Genotyping BeadChips (https://www.illumina.com). We focused on missense variants, because this variant type results in a moderate to very high phenotypic effect and induces qualitative changes in the encoded protein^25^. Seven missense variants in *ULK4*, of those displayed in the UCSC track ‘dbSNP151 in >= 1% of samples‘, were on the array:

- rs17215589 [<u>G</u>CT><u>A</u>CT] alias VAR_029009 coding for A715T
- rs3774372 [A<u>A</u>A>A<u>G</u>A] alias VAR_029006 coding for K569R
- rs1052501 [<u>G</u>CT><u>A</u>CT] alias VAR_029005 coding for A542T
- rs1716975 [<u>A</u>TT><u>G</u>TT] alias VAR_051679 coding for I224V
- rs2272007 [A<u>A</u>A>A<u>G</u>A] alias VAR_041287 coding for K39R
- rs4973986 [<u>T</u>CC><u>G</u>CC] alias VAR_029008 coding for S640A
- rs6769117 [G<u>C</u>G>G<u>T</u>G] alias VAR_051680 coding for A1261V

Of these seven missense variants in linkage disequilibrium, we determined the best variant with respect to functional signatures, on basis of the effect on gene expression and protein stability. For gene expression, we examined influence of the variants on *ULK4* transcript ENSG00000168038.10 (GTEx Release V8; dbGaP Accession phs000424.v8.p2; https://gtexportal.org/home/). For protein stability, we investigated the effect on protein aggregation tendency, using SNPeffect4.0^26-28^ (http://snpeffect.switchlab.org/).

## Results

### Variant Prioritization

Among the seven missense variants under investigation, only rs17215589 which codes for an exchange of alanine[A]-to-threonine[T] at amino acid position 715 of the ULK4 protein, showed an effect on both functional indicators, i.e., gene expression and protein aggregation tendency (Table S1, Figure S1). Therefore, we prioritized variant rs17215589 for our imaging genetics analysis. According to the Expert Protein Analysis System (ExPASy)^29^, rs17215589 changes the properties of the amino acid at position 715 of the ULK4 protein from small size and hydrophobic [A] to medium size and polar [T] (https://web.expasy.org/variant_pages/VAR_029009.html).

### Imaging Genetics Analysis

Genotype distribution of rs17215589 was N=3 AA, N=61 GA, N=170 GG, and did not deviate from Hardy-Weinberg equilibrium (P_HWE_=0.434). In order to explore the effects of *ULK4* on brain regions within the extended reward system, whole-brain group analyses were conducted comparing groups differing with respect to the rs17215589, using the contrast minor allele homozygotes + heterozygous minor allele carriers *vs*. major allele homozygotes (search criterion P<0.005, uncorrected; see Table 1 and Figure 1).

**Table 1.** Effects of *ULK4* variant rs17215589 on reward-related brain activation. Results from the contrast minor allele carriers > major allele homozygotes; L: left; R: right. Regions surviving family-wise error correction at P<0.05 using a priori coordinates from previous studies are indicated in bold.

| *Brain Region* | *MNI coordinates* | *Peak Level*<br>*T-value* | *Peak Level*<br>*P-value<sub>uncorr.</sub>* |
| --- | --- | --- | --- |
| L - ventral striatum | -18 18 3 | 3.28 | 0.001 |
| R - ventral striatum | 21 18 3 | 2.71 | 0.004 |
| **L/R - ventral tegmental area\*** | 0 -21 -9 | 3.21 | 0.001 |
| **R - ventral tegmental area\*** | 12 -24 -12 | 2.78 | 0.003 |
| L - anteroventral prefrontal cortex<sup>#</sup> | -12 33 12 | 3.83 | <0.001 |
| R - anteroventral prefrontal cortex | 33 48 0 | 2.93 | 0.002 |
| **L - middle frontal gyrus\*** | -27 39 15 | 3.81 | <0.001 |
| R - middle frontal gyrus | - | - | - |
| L - frontomedian cortex | -12 21 42 | 3.29 | 0.001 |
| R - frontomedian cortex | 12 21 36 | 3.65 | <0.001 |
| L - intraparietal cortex | -30 -51 42 | 2.62 | 0.005 |
| R - intraparietal cortex | - | - | - |
| L - inferior frontal junction | -39 6 36 | 3.82 | <0.001 |
| R - inferior frontal junction | - | - | - |
| L - fusiform gyrus | -24 -51 -3 | 3.99 | <0.001 |
| R - fusiform gyrus | - | - | - |
| L - orbitofrontal cortex | -36 39 -6 | 3.58 | <0.001 |
| R - orbitofrontal cortex | 33 39 -6 | 3.16 | 0.001 |
| L - middle temporal gyrus | -39 -18 -12 | 4.59 | <0.001 |
| R - middle temporal gyrus | - | - | - |
| L - frontal eye field<sup>#</sup> | -36 -6 57 | 3.15 | 0.001 |
| R - frontal eye field | - | - | - |

\* $P_{\text{FWE-corrected}} < 0.05$ , $^{\#}P_{\text{FWE-corrected}} < 0.1$ for small volume analysis (SVC, sphere of 9 mm radius) around a-priori coordinates from Trost et al (2014)<sup>7</sup>.

**Figure 1.**
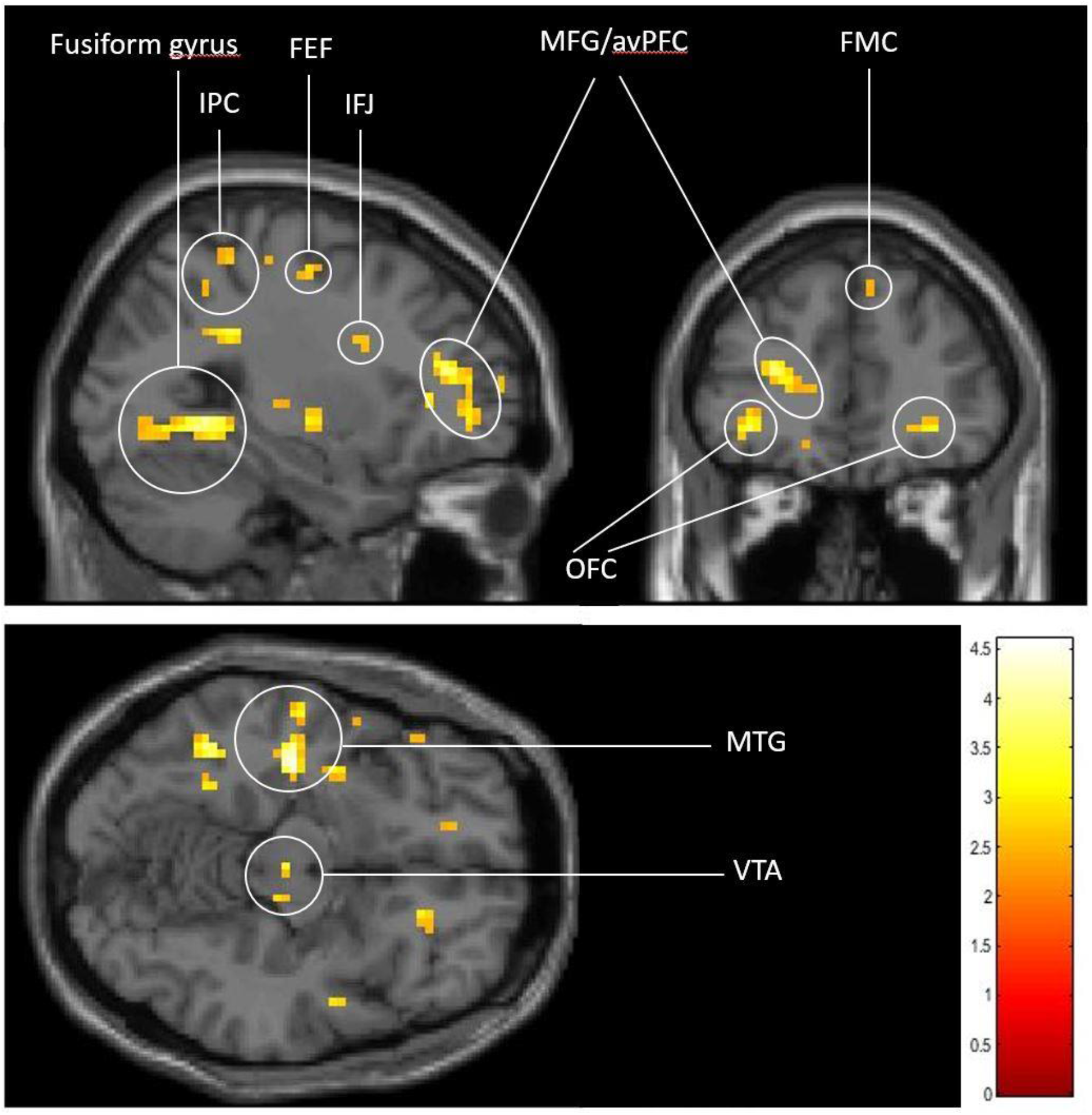
Effects of *ULK4* rs17215589 on activation of the extended reward system. For illustration purposes, the T-map of these effects is shown at a lowered threshold of p < 0.01, uncorr. and at the MNI-coordinates: -27 39 -12; FMC: frontomedian cortex; IFJ: inferior frontal junction; IPC: intraparietal cortex; MFG/avPFC: middle frontal gyrus/anteroventral prefrontal cortex; MTG: middle temporal gyrus; OFC: orbitofrontal cortex; VTA: ventral tegmental area.

The analysis demonstrated effects on reward-related activation in the left ventral striatum (MNI coordinates: -18 18 3, T-value: 3.28, P_uncorr_=0.001), right ventral striatum (MNI coordinates: 21 18 3, T-value: 2.71, P_uncorr_=0.004), left/right ventral tegmental area (MNI coordinates: 0 -21 -9, T-value: 3.21, P_uncorr_=0.001), and right ventral tegmental area (MNI coordinates: 12 -24 -12, T-value: 2.78, P_uncorr_=0.003) in terms of increased activation in rs17215589 minor allele carriers.

In addition to the effects on these subcortical core areas of reward processing, whole-brain analysis revealed a bilateral increase of activation in further brain regions of the extended reward system, e.g., in the anteroventral prefrontal cortex (left: MNI coordinates -12 33 12, T-value: 3.83, P_uncorr_<0.001; right: MNI coordinates: 33 48 0, T-value: 2.93, P_uncorr_=0.002), and in the frontomedian cortex (left: MNI coordinates: -12 21 42, T-value: 3.29, P_uncorr_=0.001; right: MNI coordinate: 12 21 36; T-value: 3.65; P_uncorr_<0.001), again in the minor allele carriers.

In the left hemisphere, we detected increased responsivity in minor allele carriers in the middle frontal gyrus (MNI coordinates: -27 39 15, T-value: 3.81, P_uncorr_<0.001), intraparietal cortex (MNI coordinates: -30 -51 42, T-value: 2.62, P_uncorr_=0.005), inferior frontal junction area (MNI coordinates: -39 6 36, T-value: 3.82, P_uncorr_<0.001), and in the fusiform gyrus (MNI coordinates: -24 -51 -3, T-value: 3.99, P_uncorr_<0.001).

Most importantly, small volume corrections (SVC) using a-priori coordinates previously reported in a sample of healthy controls^7^ confirmed the significance of these *ULK4* gene effects on reward-related brain activation in the left and right VTA and the left middle frontal gyrus (p<0.05, FWE-corrected). No significant genotype effects were observed in the opposite direction, i.e. in terms of reduced activations in rs17215589 minor allele carriers.

## Discussion

The aim of this study was to investigate the effects of genetic variation in the candidate gene *ULK4* on brain responses to conditioned reward stimuli within the mesolimbic reward system. Consistent with our expectation, functional genetic variation in *ULK4* revealed genotype effects on reward-related brain activation in several key regions of the extended reward system, among them the ventral tegmental area, the nucleus accumbens and the middle frontal gyrus.

Since the initial reports on *ULK4* and its association with psychiatric disorders, accumulating evidence suggests that *ULK4* is crucial for a variety of brain neuronal processes, including neurogenesis, neuronal motility, myelination, cilia maintenance, white matter integrity, and corticogenesis^30^. *ULK4* regulates a number of biochemical pathways, e.g., mitogen-activated protein kinase (MAPK) pathway, p38 mitogen-activated protein kinase (p38) pathway, c-Jun N-terminal kinase (JNK) pathway, and protein kinase C (PKC) signalling. One of these pathways, or a combination of them, when altered by knockdown of *ULK4*, was reported to reduce microtubule stability by diminishing alpha-tubulin acetylation^19^, an indicator for stable microtubules^31^. Intact microtubules play an important role for trafficking of key molecules of the dopaminergic neurotransmitter system, e.g. the dopamine transporter^32^.

Dopaminergic neurons are particularly vulnerable to disruption of microtubules. One study assessed the effect of microtubule depolymerization by the microtubule disruptor colchicine, on dopamine fibers in the mouse striatum and concluded that microtubule dysfunction may play a significant role in the death of dopamine neurons^33^. Another study in mice that applied nocodazole, a microtubule depolymerization reagent, showed that this substance damaged dopamine neurons and increased depression-like behavior, whereas epothilone, a microtubule-stabilizing agent, had the opposite effect^31^. These studies underline the importance of microtubule stability as regulator of dopaminergic neurotransmission.

A further possible molecular mechanism that may mediate the effects of *ULK4* on reactivity of the extended reward system is the Akt-GSK-3 pathway. In an exploration of molecular mechanisms possibly leading from *ULK4* to schizophrenia-like behavior, Hu et al (2022) found that in *ULK4* conditional knockout mice, in which this gene was deleted in the cerebral cortex and hippocampus, Akt-GSK-3 signaling was elevated. It is well known that GSK-3 is the main downstream substrate of Akt, but how exactly deficiency in *ULK4* leads to altered Akt-GSK-3 signaling remains largely unknown. Although it was suggested that *ULK4* acts on the phosphatase PP2A which works together with Akt upstream of GSK-3 and regulates its activity by balancing phosphorylation/dephosphorylation, the full molecular mechanism is unknown^34^. Nevertheless, reminiscent of the reports concerning *ULK4*^19, 22, 35^, also alterations of the GSK-3 regulatory pathway have been reported to be involved in psychiatric diseases^36, 37^ and in the response of mental disorders to drugs^38-40^.

Akt and GSK-3 are also key factors for the intracellular signalling cascade following dopamine receptor activation in vivo, and are therefore highly relevant for our study. Pharmacological dopamine receptor activation was shown to result in the modulation of the activity of Akt and GSK-3. In mice, treatment with amphetamine, an indirect agonist on dopamine receptors that acts by increasing extracellular synaptic dopamine, was demonstrated to lead to inhibition of Akt activity and activation of GSK-3. Additional evidence was generated by administration of apomorphine, a direct D1/D2 dopamine receptor agonist, which also reduced Akt activity, and confirmed that dopamine regulates the Akt-GSK-3 pathway^41^.

In conclusion, we found support for a yet unreported functional influence of *ULK4*, namely the modulation of extended reward system responses to conditioned stimuli. Thus, our study adds *ULK4* as a further gene to the few already known genetic influence factors on reactivity of the reward system in humans. Replication studies and further functional analyses are warranted to corroborate the effects of genetic variation of *ULK4* on reward system reactivity.

## Supplementary material

Supplementary material is available online.

## Acknowledgments

We thank all subjects who participated in this study. We also thank Maria Keil, Center for Translational Research in Systems Neuroscience and Psychiatry, Department of Psychiatry and Psychotherapy, Georg-August-University Göttingen, Göttingen, for support in recruitment and fMRI investigation of volunteers. We thank Prof. Dr. Elisabeth Binder and Monika Rex-Haffner, Max-Planck-Institute for Psychiatry, Munich for genome-wide SNP genotyping.

