## Supplementary material for "Impact of Unc-51 Like Kinase 4 (*ULK4*) on the Reactivity of the Extended Reward System in Response to Conditioned Stimuli"

### Methods

*Expression Quantitative Trait Loci (eQTL)*

Five of the selected seven variants, i.e., rs17215589, rs3774372, rs1052501, rs1716975, and rs2272007 showed strong allele-specific influences on the *ULK4* transcript in brain tissues in the GTEx database (https://gtexportal.org/home/), with P-values ranging from P=0.000015 to P=4.9e-33 (Table S1).

*Protein stability*

Loss of protein structure stability is a major disease-causing factor[1]. One possible molecular phenotype of an amino-acid exchange is influence on protein aggregation[2, 3]. ß-aggregation occurs when intermolecular ß-sheets are formed that are initiated by amino-acid sequence stretches which act as nuclei for ß-aggregation[4]. These amino-acid sequence stretches are short aggregation-prone regions that are buried in the folded state in the hydrophobic protein core and are not exposed to the solvent. In the core, the short-aggregation-prone regions play a role in stabilization of tertiary structure interactions of the intact protein. When the intact protein is partially unfolded, the short aggregation prone-regions become solvent-exposed, and form inter-molecular beta-strand interactions, by which proteins can self-assemble into aggregates[5].

The effect of the seven missense variants in *ULK4* on protein ß-sheet aggregation was examined using the SNPeffect4.0 tool[5] (http://snpeffect.switchlab.org/). In SNPeffect4.0, the TANGO algorithm identifies aggregation-prone regions in proteins, by analyses of the hydrophobicity and the propensity to form a beta-sheet. A TANGO region is usually buried in the protein core which means that its presence does not necessarily cause that the protein readily aggregates. However, if an amino-acid exchange that influences the stability is located in a TANGO region, the protein aggregation can become less or more prominent than in the wild type protein. For each amino acid residue in a protein, TANGO calculates the percent occupancy of ß-aggregation conformation[2].

Of the seven missense variants, only rs17215589 had a divergent value in respect protein stability compared to the wild type protein, indicating a tendency for decreased aggregation of the ULK4 protein due to rs17215589 (Figure S1 a,b).

### Supplementary Tables

**Table S1. Expression quantitative trait loci (eQTL) of *ULK4* missense variants.** Data from GTEx Release V8 (dbGaP Accession phs000424.v8.p2; https://gtexportal.org/home/); NA: not listed as eQTL.

| ***Brain region*** | ***P-value*** | | | | | | |
|---|---|---|---|---|---|---|---|
| | *rs17215589* | *rs3774372* | *rs1052501* | *rs1716975* | *rs2272007* | *rs4973986* | *rs6769117* |
| Frontal Cortex | 1.2e-23 | 1.7e-23 | 1.6e-27 | 3.1e-26 | 3.6e-28 | NA | NA |
| Cortex | 3.5e-21 | 1.1e-23 | 1.5e-31 | 8.2e-32 | 4.9e-33 | NA | NA |
| Putamen | 3.4e-21 | 2.1e-22 | 5.4e-26 | 9.8e-27 | 4.2e-28 | NA | NA |
| Cerebellum | 1.2e-21 | 4.0e-21 | 7.9e-24 | 7.2e-23 | 3.3e-24 | NA | NA |
| Anterior cingulate cortex | 4.5e-18 | 3.7e-20 | 3.0e-23 | 1.3e-19 | 3.0e-23 | 0.000029 | NA |
| Nucleus accumbens | 2.6e-18 | 3.1e-21 | 3.7e-25 | 1.4e-23 | 1.3e-25 | 0.0000068 | NA |
| Cerebellar Hemisphere | 1.2e-15 | 1.0e-15 | 6.7e-18 | 1.2e-16 | 5.8e-18 | NA | NA |
| Caudate | 3.5e-14 | 1.3e-14 | 5.8e-16 | 8.2e-16 | 1.4e-16 | NA | NA |
| Hypothalamus | 2.9e-14 | 9.7e-13 | 7.9e-14 | 5.1e-13 | 7.9e-14 | NA | NA |
| Hippocampus | 6.8e-14 | 5.3e-13 | 1.2e-14 | 1.1e-13 | 1.2e-14 | NA | NA |
| Substantia nigra | 6.9e-12 | 4.5e-10 | 3.7e-12 | 8.0e-13 | 8.0e-13 | NA | NA |
| Spinal cord | 2.7e-14 | 1.1e-12 | 9.0e-15 | 1.6e-14 | 9.0e-15 | NA | NA |
| Amygdala | 1.3e-7 | 0.000015 | 4.2e-8 | 7.5e-9 | 1.0e-8 | NA | NA |

### Supplementary figures

**a)** rs17215589 (A715T)

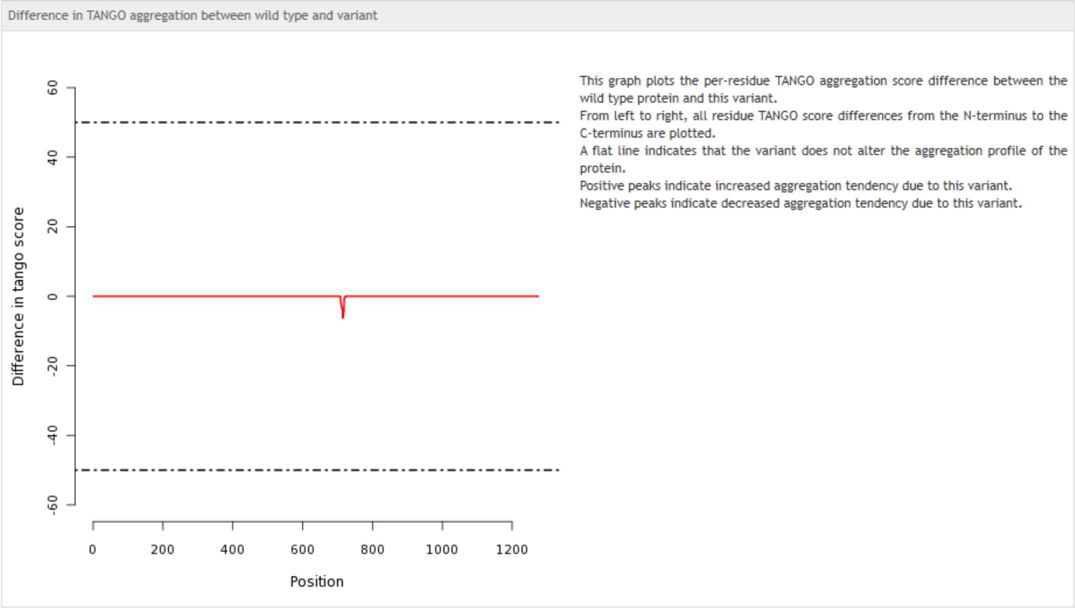


**b)** rs3774372 (K569R), rs1052501 (A542T), rs1716975 (I224V), rs2272007 (K39R), rs4973986 (S640A), rs6769117 (A1261V).

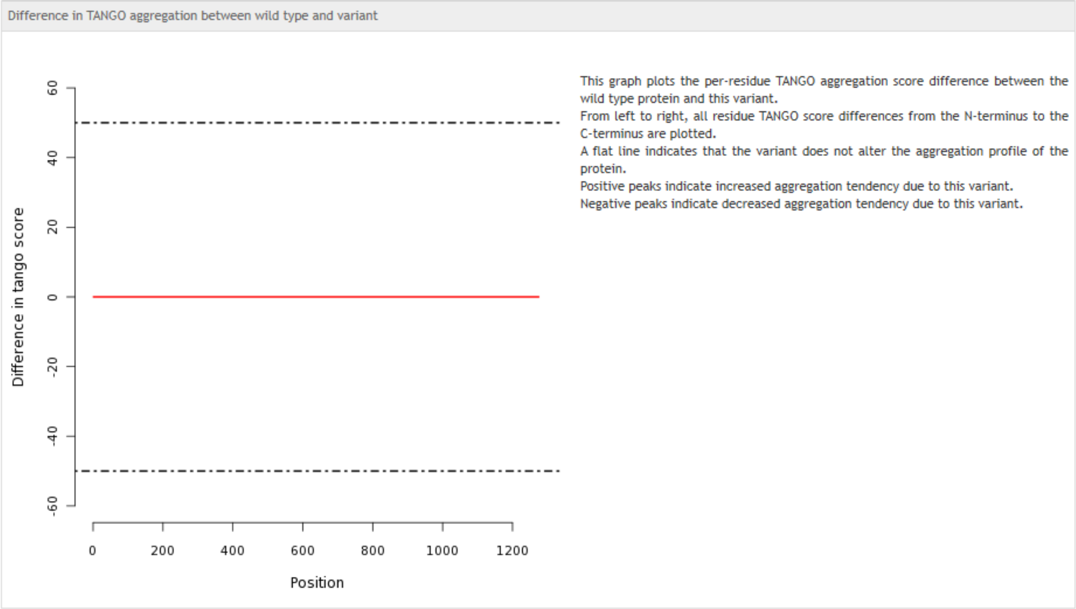


**Figure S1 a,b. Aggregation tendency of the variants**. Plots according to SNPeffect4.0 (http://snpeffect.switchlab.org). Only rs17215589 (A715T) diverges from the wild type. The negative peak for rs17215589 indicates a tendency to decreased aggregation due to this variant.
